# Neuronal firing in the medial temporal lobe reflects human working memory workload, performance and capacity

**DOI:** 10.1101/2020.06.15.152207

**Authors:** Ece Boran, Peter Hilfiker, Lennart Stieglitz, Thomas Grunwald, Johannes Sarnthein, Peter Klaver

## Abstract

2

The involvement of the medial temporal lobe (MTL) in working memory is controversially discussed. Critically, it is unclear whether and how the MTL supports performance of working memory. We recorded single neuron firing in 13 epilepsy patients while they performed a visual working memory task. The number of colored squares in the stimulus set determined the workload of the trial. We used the subjects’ memory capacity (Cowan’s K) to split them into a low and high capacity group. We found MTL neurons that showed persistent firing during the maintenance period. Firing was higher in the hippocampus for trials with correct compared to incorrect performance. Population firing predicted workload particularly during the maintenance period. Prediction accuracy of single trial activity was strongest for neurons in the entorhinal cortex of low capacity subjects. We provide evidence that low capacity subjects recruit their MTL to cope with an overload of working memory task demands.

**Significance:** Humans are highly limited in processing multiple objects over a short period of time. The capacity to retain multiple objects in working memory is typically associated with frontal and parietal lobe functioning, even though medial temporal lobe (MTL) neural architecture seems capable to process such information. However, there are conflicting findings from patient, electrophysiological and neuroimaging studies. Here we show for the first time that correct performance, workload and individual performance differences are reflected in separate mechanisms of neural activity within the MTL during maintenance of visual information in working memory. The data suggest that low capacity subjects use the MTL to process the overload of information.

## 3 Introduction

Humans need to interpret a continuous flow of information from the immediate surroundings contingently on internal goals and prior knowledge. Fortunately, in many real-life situations such as in traffic, group conversation or team sports, humans can anticipate and use prior knowledge to mitigate processing limitations. Substantial individual differences emerge when the demands to update and maintain information in working memory are high, i.e. when visuospatial information needs to be processed simultaneously (1, 2). These individual differences are relevant throughout life, associated with intelligence and are a good predictor of school aptitude, particularly for math skills (3, 4).

Updating and maintaining working memory content requires collaboration between multiple areas in the brain. Core neural structures are located in the posterior parietal cortex and prefrontal cortex, while the role of the MTL is unclear (for reviews see (5, 6)). The posterior parietal cortex supports elementary functions that are necessary for multiple object maintenance. This includes binding object features such as color and shape within the focus of spatial attention and to maintain the focus of attention to objects in space (7, 8). The prefrontal cortex is necessary for shifting control of attention between objects and updating of memory traces after interference or decay (9, 10). There is sufficient evidence from non-invasive brain stimulation studies showing the necessity of these areas in working memory, including its support by perceptual brain areas (9, 11, 12), for a review see (13).

Critically discussed is the role of the MTL and particularly the hippocampus, in working memory. Injuries to the MTL can lead to memory problems throughout development (14, 15). Yet, working memory functions were mostly spared in these patients (16, 17). The dominant hypothesis assumes that the MTL is used to retrieve memory traces from long-term memory when working memory is not operating well, suggesting that long-term memory processes complement working memory operations (18). This hypothesis is supported by findings that patients with MTL damage and long-term memory deficit failed in high working memory load conditions (many items to retain) (19). The MTL patients faced similar challenges when delay time increased (20), or when there was interference from distractors in memory (21).

Other lines of evidence are difficult to explain with the dominant hypothesis. Performance of MTL patients is impaired in two types of working memory tasks. In one type, MTL patients were impaired in performing working memory tasks with novel complex items, such as abstract objects (22) and unfamiliar faces (23, 24). This suggests an importance of encoding novel information in working memory in the MTL. Patients with MTL lesions failed also in object-location tasks, even at lower workload conditions, but not in “pure” spatial tasks or color tasks (25–27). MTL patients also made more swapping errors (28, 29), suggesting that the MTL could be involved in object-location mapping during working memory tasks. These operations might depend on the special role of the MTL in processing novel representations, including binding of items to a context during navigation (5).

Electrophysiology and neuroimaging evidence specifically links MTL activity to maintenance in working memory. A recent review on single neuron recordings suggested direct involvement of MTL neural firing patterns in working memory (6). Patterns of neural activity were consistent to specific predictions from Cowan’s activated long-term memory model of working memory. Specifically, stimulus-specific persistent neural firing was associated with online maintenance of information in working memory within the focus of attention, while a neural activity pattern that reflected specific representations of an object could reoccur after being temporarily disrupted by interfering information, which mimicked predictions of reactivated long-term memory (30, 31). BOLD fMRI studies could link MTL activity to the process of maintaining object-location information (32, 33). Two fMRI studies specifically suggested that neural activity increases with workload, but only up to the set size of the individual working memory capacity. Those studies, however, could not link the activity to maintenance processing in working memory (34, 35). It is thus unclear if neural mechanisms associated with maintaining multiple object-location bindings is useful for working memory performance.

The core aim in this study is to provide direct evidence on the role of the MTL neural firing in maintaining multiple objects in working memory under low and high workload conditions. Recent methodological developments now enable us to monitor and predict cognitive operations *in vivo* by microelectrode recording of neuronal firing in the human MTL. Studies using support vector machines on neural population firing to decode specific items reported that persistent population firing predicted memory accuracy (30, 31), while variation of working memory load could not be decoded in the hippocampus. Another study used a Sternberg delayed match-to-sample task and varied workload with the number of letters maintained in memory. In that study persistent neural firing showed workload dependent decoding accuracy during maintenance (36). Neither study reported individual differences in decoding accuracy in relation to workload and working memory capacity.

The current study recorded single neuron firing in epilepsy patients during performance of a visual working memory task. We tested for individual differences in neural activity relating to the ability to maintain multiple objects in working memory.

## 4 Results

### 4.1 Task and behavior

Subjects performed a visual working memory change detection task (16 total sessions from 13 subjects, Table S1). The task (Fig. 1a) was used in several studies using fMRI (34, 35, 37). The encoding period is temporally separated from the maintenance period. In each trial, subjects were instructed to memorize an array of 1, 2, 4 or 6 colored squares (memory array) simultaneously presented for 0.8 s (encoding). The number of squares was thus specific for the memory load. After a delay (maintenance) period of 0.9 s, a probe array of the same size prompted the subjects to indicate by button press (“Same” or “Different”) whether the array was identical to the memory array (test). This cognitively demanding task involves internal processing in the absence of external stimuli during maintenance.

**Figure 1.**
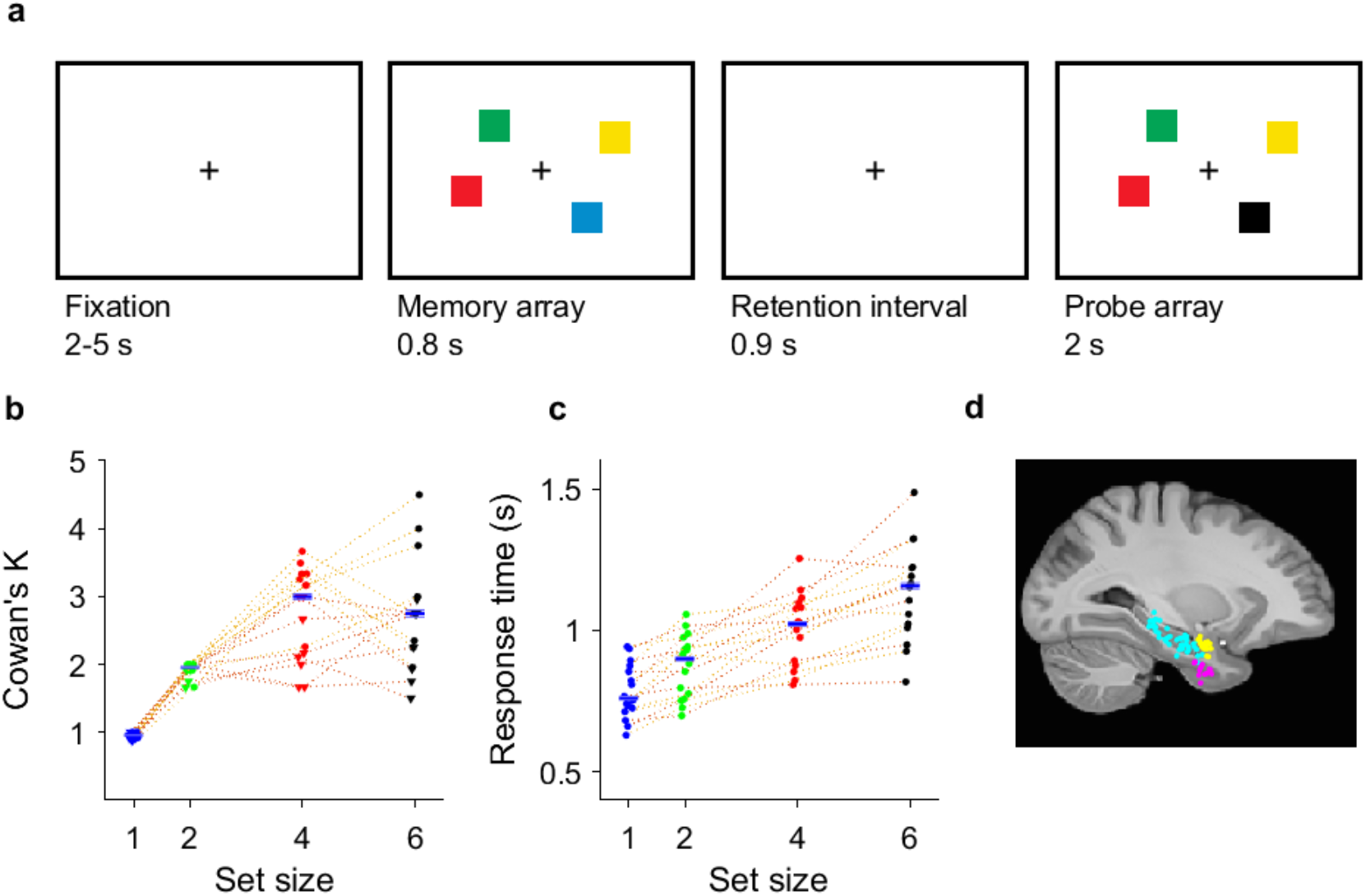
Task, behavioral results, and recording sites. **(a)** Visual working memory was examined using a change detection task. In the task, arrays of colored squares were presented and had to be memorized. The array size, called *set size*, determined WM workload (low workload: **1, 2** squares; high workload: **4, 6** squares). In each trial, presentation of a memory array (encoding period, 0.8 s) was followed by a delay (retention interval, 0.9 s). After the delay, a probe array was shown, and subjects indicated whether the probe array differed from the memory array. **(b and c)** Behavioral results. **(b)** High and low capacity subjects were separated using median memory capacity (median K_max_ = 3), where memory capacity K_max_ = max(K_ss_), Cowan’s *K* (1) for each set size (ss) = (hit rate + correct rejection rate – 1)*ss. Each dotted line connects an individual session. A total of 16 sessions were performed by 13 subjects. **(c)** Median reaction time (relative to onset of the test array) as a function of workload. The jitter reflects the rank order of the reaction time in trials with array size 6. Each dotted line connects an individual session. In **(b)** and **(c)**, orange and yellow lines represent subjects with low and high memory capacity respectively. **(d)** Location of the microelectrodes at the tip of the depth electrodes in MNI152 space. Recording locations are projected on the parasagittal plane x = −25.2 mm and are color-coded (cyan, hippocampus; magenta, entorhinal cortex; yellow, amygdala).

The subjects performed well: the rate of correct responses decreased with set size from a set size of 1 (98% correct responses) to 2 (99 %), 4 (88%) and 6 (73%) (Fig. 1b, permuted repeated-measures ANOVA, F_3,45_ = 89.79, *p* = .0002). Across all sessions, the median memory capacity was 3 (range 1.9-4.5) (Cowan’s K, (hit rate + correct rejection rate - 1) *set size), which indicates that the subjects were able to maintain at least 3 squares in memory. Separation of below and above memory capacity set sizes was in practice the same as set size separation for low and high workload. We chose the median capacity to divide subjects into a low and a high capacity group (with 7 and 6 subjects, respectively). The mean response time (RT) for the correct trials (2678 trials) increased with set size (Fig. 1c, 118 ms/item, permuted repeated-measures ANOVA, F_3,45_ = 48.47; *p* = .0002). In sum, these data show that our subjects were able to perform the task and that the difficulty of the task increased with set size.

### 4.2 Neuronal firing

To find out whether MTL neurons participated in this task, we performed multielectrode recordings of neuronal activity from microelectrodes separated into single- or multiunit activity. We refer here to a putative unit by the term “neuron”. We recorded the activity of 809 (50 ± 25 per session) neurons from the hippocampus (Hipp, n = 454), entorhinal cortex (Ent, n = 209), and amygdala (Amg, n = 146) across all microelectrodes. Low capacity subjects had 433 neurons (median (IQR) 48 (34–93) neurons per subject), high capacity subjects had 376 neurons (median (IQR) 60 (23–73) neurons per subject). We quantitatively assessed the spike sorting quality and identified the firing rate of all neurons as median (IQR) 1.09 (0.52-2.43) Hz (Supplementary Figure S1). Fig. 1d shows the microelectrode recording sites projected on a parasagittal plane in the Montreal Neurological Institute (MNI) space. In our first analysis of individual neuron types, we focused on the maintenance period, when stimuli were absent, and the test period after the presentation of the probe array. To reduce noise in the classification, unless stated otherwise, subsequent analyses included only trials with correct responses.

First, we identified neurons that fired persistently in the maintenance period. These maintenance neurons were defined by their higher firing rates during the maintenance period than during the fixation period (permutation t-test, *p* = .0005, Fig. 2a, further examples in Supplementary Figure S2). Maintenance units were found in the hippocampus, the amygdala and the entorhinal cortex (Fig. 2b).

**Figure 2.**
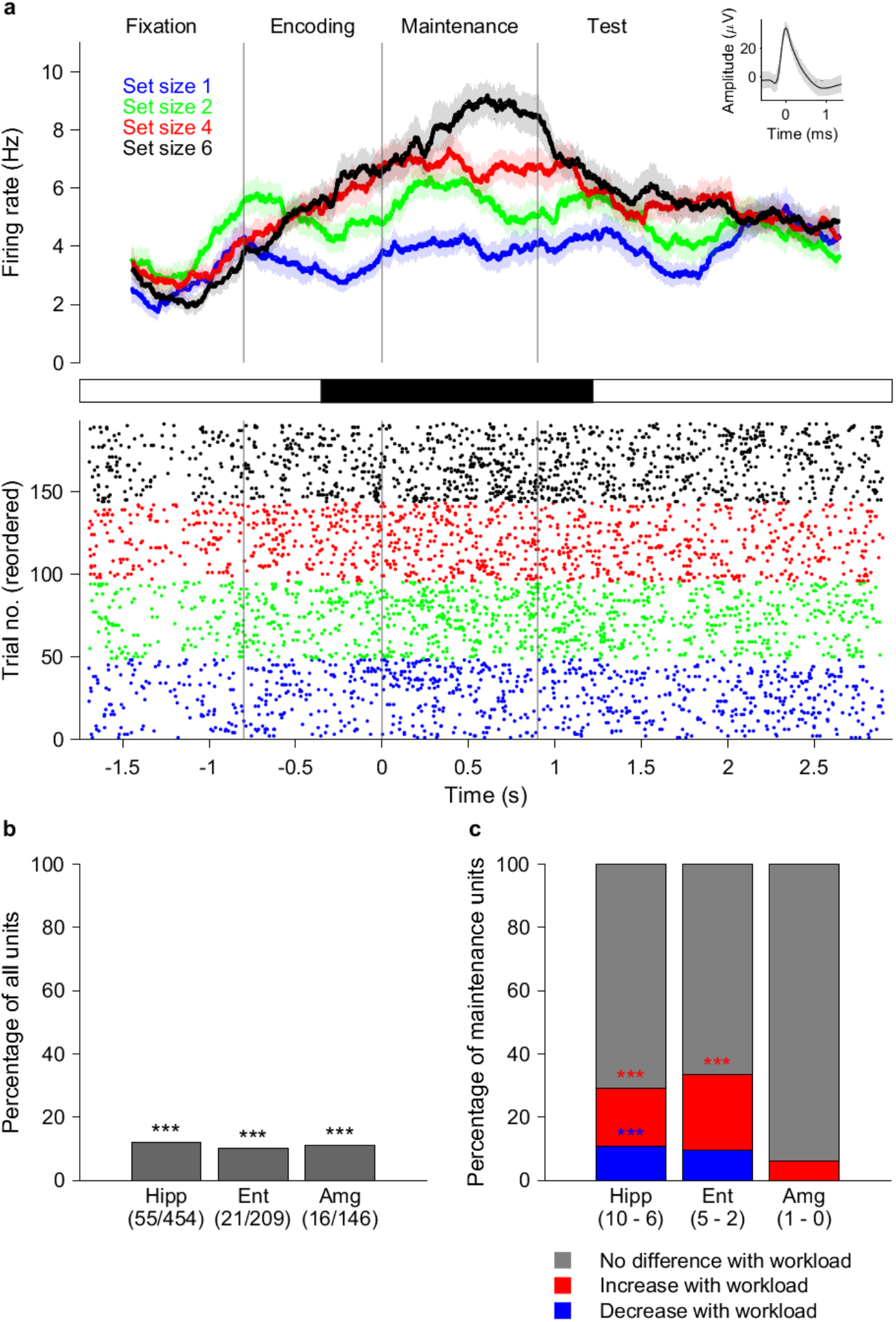
Persistent activity of maintenance neurons. **(a)** Example of a maintenance neuron recorded from entorhinal cortex. Top: Peristimulus time histogram (bin size, 500 ms; step size, 20 ms). Shaded areas represent ± SEM across trials of all spikes associated with the neuron. Inset: Mean extracellular waveform ± SEM. Middle: Periods of significance (black) between low-workload trials (set size 1 and 2, 95 trials) and high-workload trials (set sizes 4 and 6, 75 trials; *p* < .05, cluster-based nonparametric permutation test). Bottom: Raster plot of trials reordered to set size for plotting purposes only. During maintenance, compared to set sizes 1 (blue) and 2 (green), the neuron fires more for array sizes 4 (red) and 6 (black). Figure S2 provides further examples of maintenance neurons. **(b)** Percentage of all recorded neurons identified as maintenance neurons in the hippocampus (Hipp), entorhinal cortex (Ent), and amygdala (Amg). The number of maintenance neurons is provided below the area label. All MTL regions contain maintenance neurons. **(c)** Percentage of maintenance neurons for which the firing rate during maintenance differed as a function of workload. The hippocampus and entorhinal cortex stood out as containing a significant percentage of neurons that increased their firing rate under high workload. Significance was assessed by comparing with a null distribution with permuted labels. \*\*\**p* = .002.

When comparing high load vs. low load trials, we found a significant number of maintenance neurons that showed an increase in firing with workload in the hippocampus (10/55, permutation test against scrambled labels, *p* = .002, Fig. 2c) and entorhinal cortex (5/21, permutation test against scrambled labels, *p* = .002). We also found a significant number of maintenance neurons that showed a decrease in firing rate with workload in the hippocampus (6/55, permutation test against scrambled labels, *p* = .002).

When comparing correct and incorrect trials, we noted that correct performance was associated with a larger number of neurons that showed enhanced persistent firing during maintenance, i.e. that were classified as maintenance neurons. There were more of these hippocampal neurons in correct trials vs incorrect trials (55/454 vs 30/454, permutation test against scrambled labels, *p* = .002). In a subgroup analysis, this effect was significant in low capacity subjects (39/248 vs 19/248, permutation test against scrambled labels, *p* = .002), but not in high capacity subjects (16/206 vs 11/206, permutation test against scrambled labels, *p* = .1980). In line with that, there were more maintenance units for correct trials in low capacity subjects compared to high capacity subjects (39/248 vs 16/206, chi2 = 6.69, *p* = .0097). This points to the importance of the hippocampus for correct performance, particularly in low capacity subjects. If the effect would be caused by the presence of errors, the effect should be biased towards high workload trials where errors occurred. The results show that this was not the case: there were fewer hippocampal maintenance units in correct trials compared to all trials when considering only high workload trials (55/454 vs 37/454, permutation test against scrambled labels, *p* = .002). By contrast, the number of maintenance units in incorrect trials compared to all trials was similar when only considering high load trials (30/454 vs 33/454, permutation test against scrambled labels, *p* = .3060). This indicates that persistently firing hippocampal neurons during maintenance support correct performance in low capacity subjects.

Next, we identified neurons, which fired specifically after the presentation of the probe array that initiated the test period. These probe neurons were defined based on increased firing only during the presentation of the probe array relative to encoding and maintenance (permutation t-test, Fig. 3a). With a rate of 5.7%, these neurons were rare (Fig. 3b).

**Figure 3.**
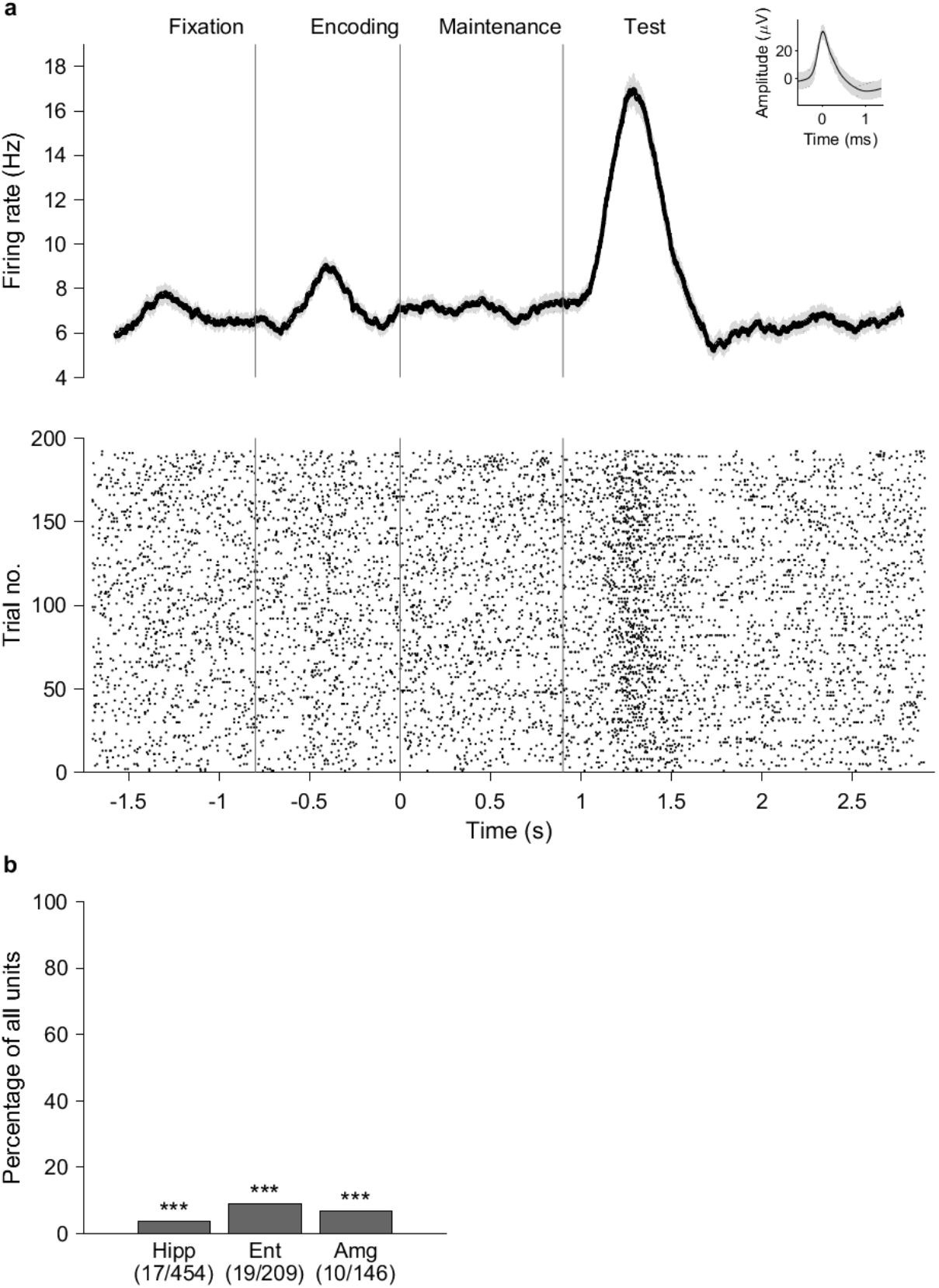
The activity of test neurons is related to working memory retrieval. **(a)** Example test neuron recorded from the entorhinal cortex. **(b)** Percentage of all recorded neurons identified as test neurons in the hippocampus (Hipp), entorhinal cortex (Ent), and amygdala (Amg). The number of test neurons is provided below the area label. All MTL regions contain test neurons. For (b), significance was assessed by comparing with a null distribution with permuted labels. \*\*\**p* = .002.

### 4.3 Neuronal population analysis

We subsequently focused on the neural population firing rate during the periods of the trial. We used demixed principal component analysis (dPCA) (38) to project the firing rates from all neurons onto a low-dimensional component space. To demix the effect of stimulus category, dPCA was informed by the workload of the trials. dPCA clearly distinguished between set sizes 1, 2, 4, and 6 (Supplementary Figure S3). For better visualization we formed a three-dimensional space from three demixed principal components (dPCs 2, 3 and 4, explaining 36% of the variance, permutation test against scrambled data, *p* = .002, Fig. 4a). The population activity distinguished between set sizes very early during the trial as seen from the projection on the dPC3-dPC4-plane, which shows four angles of 90° each, i.e. the optimal balanced distinction between the four set sizes.

**Figure 4.**
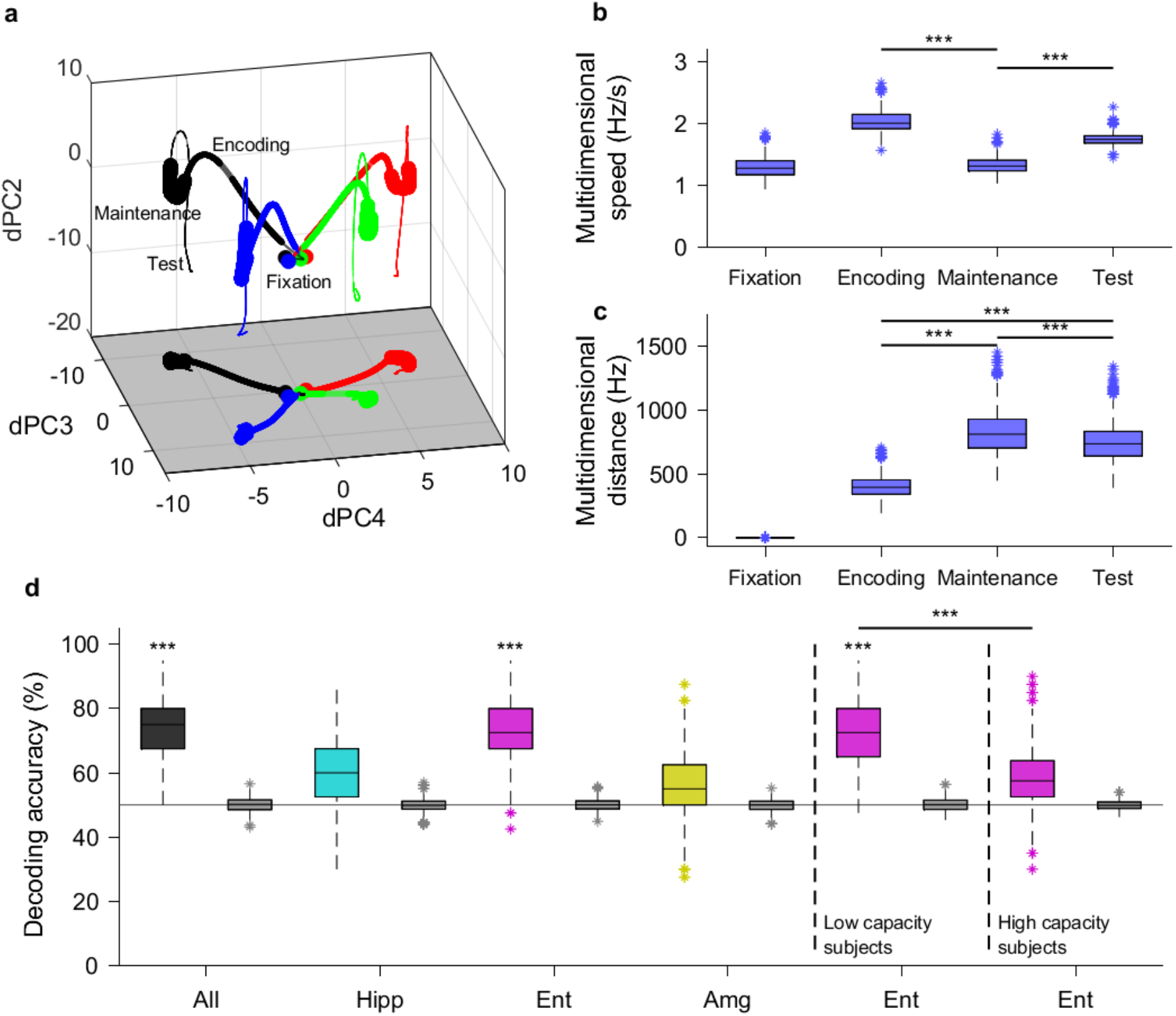
Population firing predicts workload. **(a)** Mean trajectories in the neuronal state space constituted by three dPCs during fixation (starting at the origin), encoding, maintenance, and retrieval. Set sizes are color-coded (1: blue; 2: green; 4: red; 6: black). **(b)** Multidimensional speed of the population in the four periods of a trial. The speed during maintenance was slowest. **(c)** Multidimensional pairwise distance between all possible pairs of attractors during the different periods of the task. The distance during maintenance was largest. The combination of low speed and large mutual distance provides evidence for attractors in state space during maintenance. **(d)** During maintenance, workload (set sizes 1 and 2 vs. set sizes 4 and 6) could be decoded from neurons in entorhinal cortex but not hippocampus and amygdala. Workload decoding achieved higher decoding accuracy for the subpopulation of neurons from low capacity subjects compared to high capacity subjects (*p* = .0005, permutation t-test). The analysis in **(b)** and **(c)** included the first 15 dPCs that explain 81.4% of the signal variance. Significance was assessed by permutation t test. \*\*\**p* = .0005. Boxplots represent quartiles (25%, 75%); horizontal lines represent medians; whiskers show ranges up to 1.5 times the interquartile range; dots outside whiskers show outliers. Markers below bars indicate significance versus chance performance. Significance was assessed on the basis of a null distribution with scrambled labels (gray boxes). \*\*\**p* = .001.

In this dPCA neural state space, we included the first 15 dPCs that explain 81.4% of the signal variance to analyze the rate of change over time (speed) (Fig. 4b). Multidimensional speed was highest during encoding, when maintenance neurons ramped up their firing. Speed was lowest during maintenance (permutation t-test against encoding and test, *p* = .0005). Analyzing the pairwise distance between trajectories that corresponded to different set sizes, we found that the distances during maintenance were larger than during encoding (permutation t-test, *p* = .0005, Fig. 4c). Thus, during maintenance, the observed pattern of the firing rates of all recorded neurons resembled an attractor as a location in state space. Over the trial, the neural trajectories clearly distinguished the periods within trials, among which the maintenance period stood out in clustering neural activity around attractors. More importantly, MTL population activity clearly distinguished between trials of set size 1, 2, 4, and 6 and was thus indicative of workload.

### 4.4 Decoding of information during the trial

We next asked, whether the neuronal activity was related to the individual subject’s working memory capacity. We analyzed the firing of each subject’s neuronal population on a trial-by-trial basis. Furthermore, we selected subpopulations that consisted of, e.g., only hippocampal maintenance neurons. We trained a decoder on a subset of trials and tested its performance on an independent set of test trials (39). This assessed whether neuronal firing was sufficiently salient to be representative of task demands or performance in single trials. Given the substantial variability in neuronal dynamics during a trial (Fig. 4a-c), we focused on the maintenance period.

For the maintenance period, we found that the workload (set size 1, 2 vs. set sizes 4, 6) of each trial could be decoded with median decoding accuracy of 75% when all neurons were pooled. (Fig. 4d, permutation test against scrambled data, *p* = .001). In a leave-one-out analysis, the exclusion of any subject resulted in decoding accuracy in the IQR [64%-75%], which verified that the accuracy was supported by all subjects. When pooling neurons from the three anatomical regions separately, decoding accuracy was significant only for entorhinal cortex neurons (Fig. 4d column 5), but not for neurons in the other areas. For the entorhinal cortex units, the workload of each trial could be decoded by units from only low capacity subjects (median decoding accuracy 73%, permutation test against scrambled data, *p* = .001, Fig. 4d column 9), but not by units from high capacity subjects (median decoding accuracy 58%, permutation test against scrambled data, *p* > 0.05). For low capacity subjects, the prediction accuracy of entorhinal neurons was significantly better than for high capacity subjects (Fig. 4d column 9 and 11, permutation t-test, *p* = .0005). This points to a distinct recruitment of neural activity in low capacity subjects while they cope with high load on working memory.

## 5 Discussion

We found changes in MTL neural firing patterns across neurons in the MTL and particularly in the entorhinal cortex and hippocampus. These changes depended on workload (the number of items in the array), capacity (whether the subject can correctly detect changes from a small or large array of items) and performance accuracy (whether the subject responds correctly to a change or no change).

### 5.1 MTL supports object separation in working memory

All four workload conditions could be separated by neural population firing in the MTL, particularly during maintenance, where the components were stable and distinguishable. Such a mechanism would allow for a reduction of noise between separate items in memory and therefore allow for a reduced chance of errors. This would fit with the hypotheses that the MTL can support object separation in working memory (28, 29, 40). The results confirm and extend on the report that showed workload dependent separation of neural firing patterns during maintenance of letters in working memory (36), but contrasts with other findings in which separation of neural firing patterns during maintenance did not predict load on complex stimuli in working memory (30, 31). This suggests that maintaining multiple simple items, including objects and letters, in working memory can be characterised by separation of neural population firing.

### 5.2 MTL is activated only in low capacity subjects

Workload during maintenance could further be predicted by the trial-to-trial variability of neural firing patterns in the MTL. This was particularly the case in the entorhinal cortex, and specifically in subjects with low working memory capacity. The entorhinal cortex is a gateway to the hippocampus receiving information from the neocortex and directing input via the perforant path to the dental gyrus of the hippocampus and is thus the station where novel information is processed first in working memory (41). Persistent firing and theta oscillations and the entorhinal cortex and hippocampus have been suggested to support processing novel items in working memory (42, 43). This is achieved by means of phase locking high frequency neural firing to theta oscillations (40, 44, 45). Our earlier report showed that verbal workload within capacity limits could be predicted by trial-to-trial variability of neural firing in the hippocampus (36). The current study used memory arrays of objects that were designed to exceed working memory capacity. We suggest that good prediction of workload in the entorhinal cortex in low capacity subjects reflects processing an overload of information and continued memorization of the items during maintenance. High capacity subjects seem more efficient in memorization, so that the entorhinal cortex is less involved during working memory maintenance.

### 5.3 Correct performance associated with increased hippocampal neural firing

A small portion of neurons in the hippocampus and entorhinal cortex increased firing during maintenance of objects. Importantly, firing rate was higher for correct than for incorrect trials by maintenance neurons, particularly in the hippocampus. We only found this effect in low capacity subjects. This is a novel finding and shows that persistent hippocampal neural firing is supportive of visual working memory performance, particularly in low capacity subjects.

### 5.4 Interpreting persistent firing

We suggest that in low capacity subjects, entorhinal and hippocampal persistent firing is instrumental to maintain multiple objects in memory and to achieve correct performance. Persistent posterior parietal activity is known to support performance at high workload in high capacity subjects, but not in low capacity subjects (3, 46, 47). It is thus tenable that low capacity subjects might use entorhinal and hippocampal activity to compensate for impaired posterior parietal functioning. The findings extend to a BOLD fMRI study with the same task showing a similar pattern for low capacity subjects, except that high capacity subjects further increased activity at high workload and that BOLD fMRI activity could not be univocally be identified as maintenance activity (35). Further, our findings add to other single cell recording studies. A verbal working memory task reported increased firing in a substantial portion (~20%) of maintenance units with workload but performance could not be predicted (36), which might be related to the lack of need to bind between object features (color-location vs. letters) and associated higher capacity for verbal items. Other studies showed that certain cells, so called concept cells, showed preferred neural activity for specific complex pictures and that persistent neural activity of concept cells predict correct behavior of a trial (30, 31). In contrast to those studies we found trial-to-trial variability of neural activity predicting workload in the entorhinal cortex, and elevation of neural firing for correct trials in the hippocampus, while both these effects were found particularly in low capacity subjects. This suggests that persistent activity is a relevant marker not only for correct maintenance of items that show category selective responses but also for workload and correct performance during memorization of arbitrary arrays of simple objects. Both manifestations of persistent activity reflected different cognitive phenomena and in different neural structures, suggesting a separation of working memory operations within the MTL.

### 5.5 Conclusions

Our findings elucidate the role of persistent firing in MTL during human working memory maintenance. We propose that subjects with low working memory capacity recruit the persistent firing in their entorhinal cortex to cope with working memory load and that the persistent firing in their hippocampus supports correct performance.

## 6 Methods

### 6.1 Subjects

Subjects were patients with epilepsy undergoing pre-operative monitoring with implanted electrodes in the MTL (13 subjects, age 18-56 years, 7 male, Table S1). All subjects had normal or corrected-to-normal vision and were right-handed as confirmed by neurophysiological testing.

### 6.2 Task

Visual working memory was examined using a change detection task (2, 35), where arrays of colored squares were presented and had to be memorized (Fig. 1a). The set size (low workload: 1, 2 squares high workload: 4, 6 squares, total 192 trials per session) determined working memory workload. In each trial, presentation of a memory array (encoding period, 0.8 s) was followed by a delay (retention interval, 0.9 s). After the delay, a probe array was shown, and subjects indicated whether the probe array differed from the memory array. A subject’s working memory capacity was estimated using Cowan’s *K (K = (hit rate + correct rejection rate – 1)*N*, where *N* = the number of objects presented (1)). High and low capacity subjects were separated on the basis of median split for Cowan’s *K*_max_ (Fig. 1b).

### 6.3 Recordings and analysis

We recorded with microelectrodes in the patients’ medial temporal lobe. Spike sorting was done using Combinato (48). Neurons active during multiple sessions entered the analysis independently for each session. Fig. 1d shows the recording sites. We first identified units that show higher activity during maintenance compared to fixation using permutation tests against scrambled labels. These were termed maintenance units. We also identified test units, which are units that show higher activity during test compared to maintenance.

### 6.4 Multidimensional state space

To illustrate the firing rates of all neurons over the periods of a trial, we used dPCA as a dimensionality reduction method (38). In contrast to PCA, which explains the variance of components regardless of task conditions, dPCA incorporates the task conditions, i.e., set size in our task, when explaining the variance. We used non-overlapping bins of 2 ms for neuronal firing and smoothed the rates in a window of 1000 ms with a Gaussian kernel. We z-scored the resulting rates based on the mean of the firing rate during fixation. We computed time-dependent and set size-dependent dPCA components and ordered them by explained variance. To create a distribution for multidimensional speed in different trial phases, we ran dPCA analysis 500 times, subsampling (20%, with replacement) neural firing rate trajectories for each run.

### 6.5 Decoding of information at the trial level

We analyzed neuronal firing at the population level on a trial-by-trial basis. We grouped the neurons according to their anatomical region (hippocampus, entorhinal cortex or amygdala). We collected the rates of all neurons and all trials (1 ms bin, 1000 ms window). We trained and tested the classifier at every 100 ms using a support vector machine (SVM) classifier, as implemented in the NDT toolbox (39). We performed 500 resampling runs using 20 cross-validation splits. We tested significance by comparing true accuracy to a null distribution of accuracies with 1000 runs of the decoder with scrambled labels. This approach resulted in a minimum p-value of .001. We corrected the *P* value for multiple comparisons using Bonferroni’s method. For the 12 tests performed (three subject groups (all subjects, low capacity, high capacity) in four regions (all, hippocampus, entorhinal cortex, amygdala), the corrected value was *P* = 0.001 × 12 = 0.012 and remained statistically significant.

## 8 Acknowledgements

We thank the physicians and the staff at Schweizerische Epilepsie-Klinik for their assistance and the patients for their participation. Funding: We acknowledge grants awarded by the Swiss National Science Foundation (SNSF 320030_176222 to J.S.), Mach-Gaensslen Stiftung (to J.S.), Stiftung für wissenschaftliche Forschung an der Universität Zürich (to J.S.). The funders had no role in the design or analysis of the study.

## Ethics statement

The study was approved by the IRB, and all subjects gave written informed consent.

## Author contributions

P.K. designed the task; P.H. and J.S. set up data acquisition; P.K, J.S. collected data; L.S. and T.G. treated patients; E.B., J.S. and P.K. analyzed data; E.B., P.K., J.S. wrote the paper with contributions from all authors.

## Data availability

Data will be made available after acceptance of the manuscript.

## References

1. N. Cowan, The magical number 4 in short-term memory: a reconsideration of mental storage capacity. The Behavioral and brain sciences 24, 87–114; discussion 114-185 (2001).

2. S. J. Luck, E. K. Vogel, The capacity of visual working memory for features and conjunctions. Nature 390, 279–281 (1997).

3. K. Fukuda, E. K. Vogel, U. Mayr, E. Awh, Quantity, not quality: the relationship between fluid intelligence and working memory capacity. Psychonomic Bulletin & Review 17, 673–679 (2010).

4. S. E. Gathercole, S. J. Pickering, C. Knight, Z. Stegmann, Working memory skills and educational attainment: evidence from national curriculum assessments at 7 and 14 years of age. Applied Cognitive Psychology 18, 1–16 (2004).

5. J. Jonides et al., The mind and brain of short-term memory. Annu Rev Psychol 59, 193–224 (2008).

6. J. Kamiński, U. Rutishauser, Between persistently active and activity-silent frameworks: novel vistas on the cellular basis of working memory. Annals of the New York Academy of Sciences 10.1111/nyas.14213 (2019).

7. N. Hakim, K. C. S. Adam, E. Gunseli, E. Awh, E. K. Vogel, Dissecting the Neural Focus of Attention Reveals Distinct Processes for Spatial Attention and Object-Based Storage in Visual Working Memory. Psychol Sci 30, 526–540 (2019).

8. S. L. Sheremata, D. C. Somers, S. Shomstein, Visual Short-Term Memory Activity in Parietal Lobe Reflects Cognitive Processes beyond Attentional Selection. J Neurosci 38, 1511–1519 (2018).

9. N. S. Rose et al., Reactivation of latent working memories with transcranial magnetic stimulation. Science 354, 1136–1139 (2016).

10. S. E. Cavanagh, J. P. Towers, J. D. Wallis, L. T. Hunt, S. W. Kennerley, Reconciling persistent and dynamic hypotheses of working memory coding in prefrontal cortex. Nat Commun 9, 3498 (2018).

11. H. M. Morgan, M. C. Jackson, M. G. van Koningsbruggen, K. L. Shapiro, D. E. Linden, Frontal and parietal theta burst TMS impairs working memory for visual-spatial conjunctions. Brain Stimul 6, 122–129 (2013).

12. S. Hilbert et al., Right hemisphere occipital rTMS impairs working memory in visualizers but not in verbalizers. Sci Rep 9, 6307 (2019).

13. C. H. Juan, P. Tseng, T. Y. Hsu, Elucidating and Modulating the Neural Correlates of Visuospatial Working Memory via Noninvasive Brain Stimulation. Curr Dir Psychol Sci 26, 165–173 (2017).

14. L. Squire, S. Zola-Morgan, The medial temporal lobe memory system. Science 253, 1380–1386 (1991).

15. F. Vargha-Khadem et al., Differential effects of early hippocampal pathology on episodic and semantic memory. Science 277, 376–380 (1997).

16. A. Baddeley, C. Jarrold, F. Vargha-Khadem, Working Memory and the Hippocampus. Journal of Cognitive Neuroscience 23, 3855–3861 (2011).

17. F. Vargha-Khadem, Differential Effects of Early Hippocampal Pathology on Episodic and Semantic Memory. Science 277, 376–380 (1997).

18. A. Jeneson, L. R. Squire, Working memory, long-term memory, and medial temporal lobe function. Learning & Memory 19, 15–25 (2011).

19. A. Jeneson, J. T. Wixted, R. O. Hopkins, L. R. Squire, Visual Working Memory Capacity and the Medial Temporal Lobe. Journal of Neuroscience 32, 3584–3589 (2012).

20. A. Jeneson, K. N. Mauldin, R. O. Hopkins, L. R. Squire, The role of the hippocampus in retaining relational information across short delays: The importance of memory load. Learning & Memory 18, 301–305 (2011).

21. Y. Shrager, D. A. Levy, R. O. Hopkins, L. R. Squire, Working Memory and the Organization of Brain Systems. Journal of Neuroscience 28, 4818–4822 (2008).

22. J. S. Holdstock, C. Shaw, J. P. Aggleton, The performance of amnesic subjects on tests of delayed matching-to-sample and delayed matching-to-position. Neuropsychologia 33, 1583–1596 (1995).

23. Y. Ezzyat, I. R. Olson, The medial temporal lobe and visual working memory: Comparisons across tasks, delays, and visual similarity. Cognitive, Affective, & Behavioral Neuroscience 8, 32–40 (2008).

24. E. A. Nichols, Y.-C. Kao, M. Verfaellie, J. D. E. Gabrieli, Working memory and long-term memory for faces: Evidence from fMRI and global amnesia for involvement of the medial temporal lobes. Hippocampus 16, 604–616 (2006).

25. A. M. Owen, B. J. Sahakian, J. Semple, C. E. Polkey, T. W. Robbins, Visuo-spatial short-term recognition memory and learning after temporal lobe excisions, frontal lobe excisions or amygdalo-hippocampectomy in man. Neuropsychologia 33, 1–24 (1995).

26. I. R. Olson, K. S. Moore, M. Stark, A. Chatterjee, Visual Working Memory Is Impaired when the Medial Temporal Lobe Is Damaged. Journal of Cognitive Neuroscience 18, 1087–1097 (2006).

27. C. Finke et al., The human hippocampal formation mediates short-term memory of colour–location associations. Neuropsychologia 46, 614–623 (2008).

28. Y. Pertzov et al., Binding deficits in memory following medial temporal lobe damage in patients with voltage-gated potassium channel complex antibody-associated limbic encephalitis. 136, 2474–2485 (2013).

29. P. D. Watson, J. L. Voss, D. E. Warren, D. Tranel, N. J. Cohen, Spatial reconstruction by patients with hippocampal damage is dominated by relational memory errors. 23, 570–580 (2013).

30. J. Kamiński et al., Persistently active neurons in human medial frontal and medial temporal lobe support working memory. 10.1038/nn.4509 (2017).

31. S. Kornblith, R. Quian Quiroga, C. Koch, I. Fried, F. Mormann, Persistent Single-Neuron Activity during Working Memory in the Human Medial Temporal Lobe. Current Biology 27, 1026–1032 (2017).

32. C. Piekema, R. P. Kessels, R. B. Mars, K. M. Petersson, G. Fernandez, The right hippocampus participates in short-term memory maintenance of object-location associations. Neuroimage 33, 374–382 (2006).

33. C. Ranganath, M. D’Esposito, Medial temporal lobe activity associated with active maintenance of novel information. Neuron 31, 865–873 (2001).

34. D. Y. von Allmen, K. Wurmitzer, P. Klaver, Hippocampal and posterior parietal contributions to developmental increases in visual short-term memory capacity. Cortex 59, 95–102 (2014).

35. D. Y. von Allmen, K. Wurmitzer, E. Martin, P. Klaver, Neural activity in the hippocampus predicts individual visual short-term memory capacity. Hippocampus 23, 606–615 (2013).

36. E. Boran et al., Persistent hippocampal neural firing and hippocampal-cortical coupling predict verbal working memory load. Science Advances 5 (2019).

37. P. Klaver, W. Knirsch, K. Wurmitzer, D. Y. von Allmen, Children and Adolescents Show Altered Visual Working Memory Related Brain Activity More Than One Decade After Arterial Switch Operation for D-Transposition of the Great Arteries. Dev Neuropsychol 41, 261–267 (2016).

38. D. Kobak et al., Demixed principal component analysis of neural population data. Elife 5 (2016).

39. E. M. Meyers, The neural decoding toolbox. Front Neuroinform 7, 8 (2013).

40. J. E. Lisman, O. Jensen, The theta-gamma neural code. Neuron 77, 1002–1016 (2013).

41. M. E. Hasselmo, C. E. Stern, Mechanisms underlying working memory for novel information. Trends Cogn Sci 10, 487–493 (2006).

42. E. Fransén, Functional role of entorhinal cortex in working memory processing. Neural Netw 18, 1141–1149 (2005).

43. S. Raghavachari et al., Gating of human theta oscillations by a working memory task. J Neurosci 21, 3175–3183 (2001).

44. M. E. Hasselmo, Grid cell mechanisms and function: contributions of entorhinal persistent spiking and phase resetting. Hippocampus 18, 1213–1229 (2008).

45. M. Leszczynski, J. Fell, N. Axmacher, Rhythmic Working Memory Activation in the Human Hippocampus. Cell Rep 13, 1272–1282 (2015).

46. F. Edin et al., Mechanism for top-down control of working memory capacity. Proc Natl Acad Sci U S A 106, 6802–6807 (2009).

47. E. K. Vogel, M. G. Machizawa, Neural activity predicts individual differences in visual working memory capacity. Nature 428, 748–751 (2004).

48. J. Niediek, J. Bostrom, C. E. Elger, F. Mormann, Reliable Analysis of Single-Unit Recordings from the Human Brain under Noisy Conditions: Tracking Neurons over Hours. PLoS One 11, e0166598 (2016).

