## Supplementary material for "Neuronal firing in the medial temporal lobe reflects human working memory workload, performance and capacity": Figures S1 S2 S3 Table S1

**This PDF file includes:**

Figures S1 to S3
Table S1

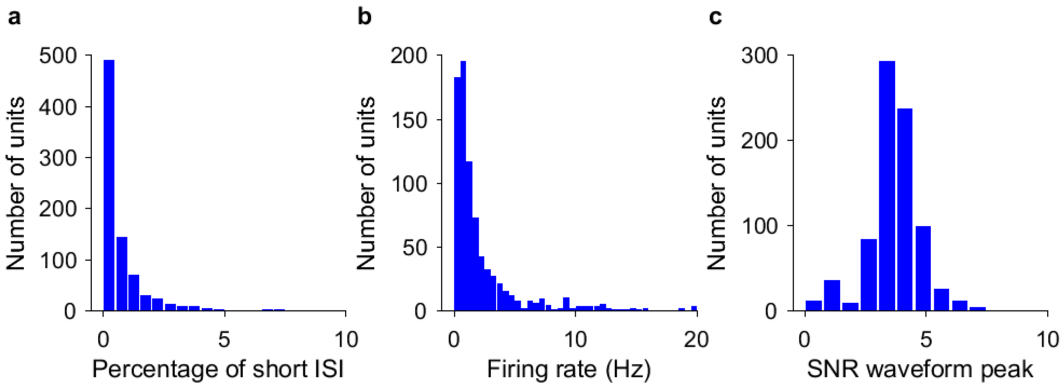


**Fig. S1. Metrics for spike sorting for the identification of putative neuronal units.**

**(a)** Histogram of percentage of inter-spike intervals (ISI) < 3 ms. The majority of units had less than 0.5% of short ISI.

**(b)** Histogram of average firing rate for all units.

**(c)** Histogram of the signal-to-noise ratio (SNR) of the peak of the mean waveform.

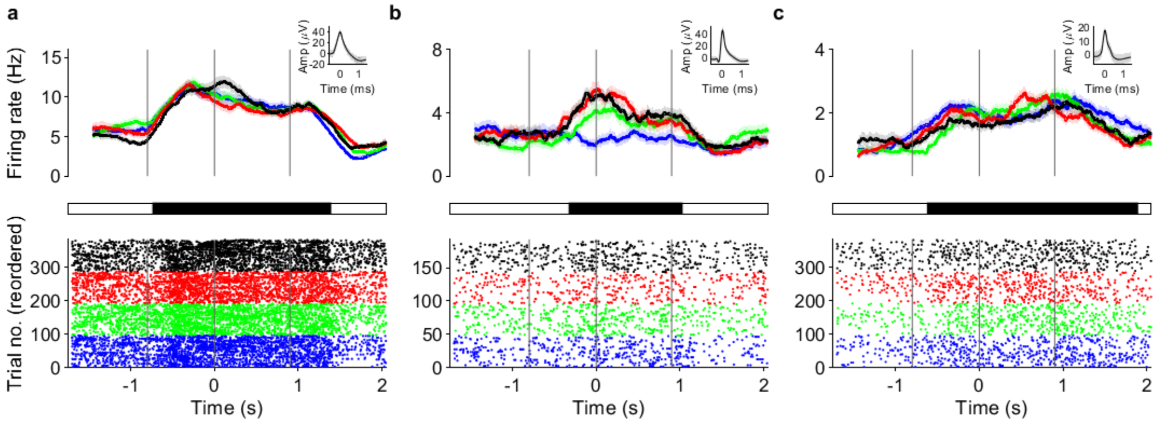


**Fig. S2.** **Examples of maintenance neurons.**

(a) An entorhinal cortex neuron from subject 3 (low memory capacity). (b) Hippocampus neurons from subject 3 and subject 8 (high memory capacity), respectively. For (a) and (c), the neuron is recorded from two sessions performed consecutively. Black bar: In these MTL neurons, the firing rate during maintenance (0 to 0.9 s) exceeds the firing rate during fixation (-1.7 to -0.8 s, cluster based non-parametric permutation test, $p < .05$). Set sizes are color-coded (1: blue; 2: green; 4: red; 6: black).

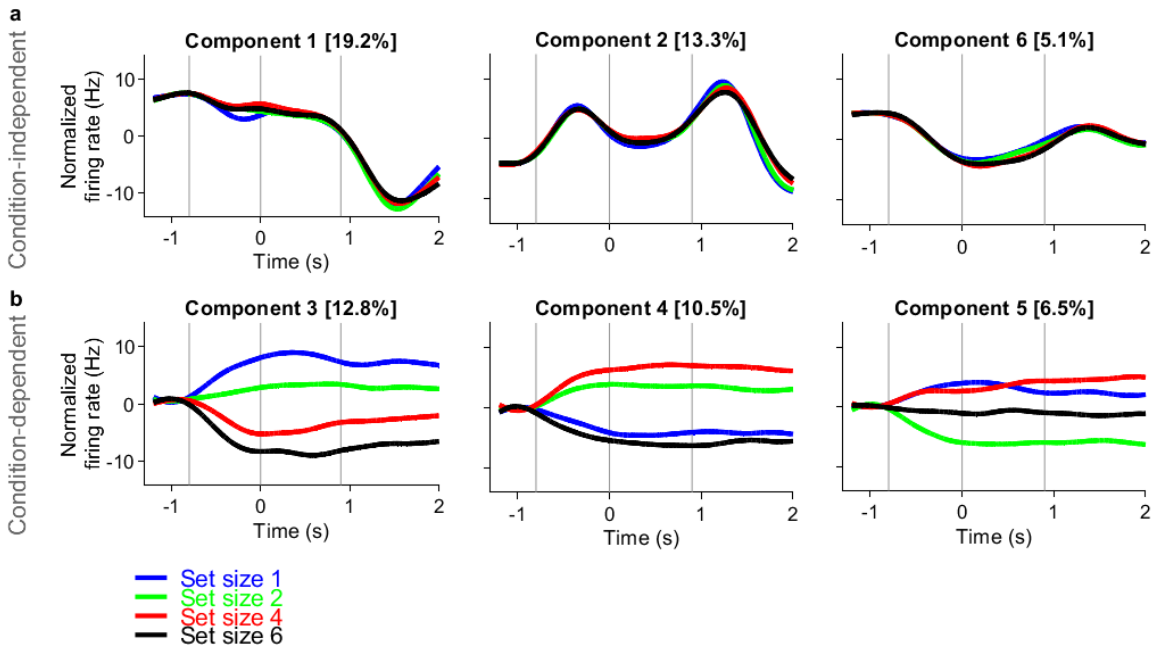


**Fig. S3. Demixed principal component analysis.**

(a) dPCA components that were modulated as a function of time independent of set size. (b) dPCA components that were modulated by set size. Components dPC3 and dPC4 together discriminate between set sizes in 2D. Set sizes are color-coded (1: blue; 2: green; 4: red; 6: black).

| Subject | Age | Sex | Pathology | Handedness | Implanted electrodes | SOZ electrodes | Number of sessions | Memory capacity (Kmax) | Number of neurons |
|---|---|---|---|---|---|---|---|---|---|
| 1 | 31 | Male | Hippocampal sclerosis | Right | AHL,AHR,AL,AR,ECL,ECR,PHL,PHR | AHL,PHL,ECL,AL | 1 | 2.11 | 33 |
| 2 | 31 | Female | Hippocampal sclerosis | Right | AHR,AR,ECR,PHR | PHR,AHR | 1 | 2.25 | 36 |
| 3 | 56 | Female | Hippocampal sclerosis | Right | AHL,AHR,AL,AR,ECL,ECR,PHL,PHR | ECR | 2 | 2.44 | 106 |
| 4 | 28 | Male | Brain contusion | Right | AHL,AHR,AL,AR,ECL,ECR,PHL,PHR | AHL,AHR,PHL,PHR | 1 | 2.75 | 85 |
| 5 | 20 | Female | Focal cortical dysplasia | Right | AHL,AHR,PHL,PHR, LL, PL | LL,PL | 1 | 2.75 | 29 |
| 6 | 35 | Male | Unknown | Right | AHL,AHR,AIR,AL,AR,ECL,ECR,PHR | AHL,AIR | 1 | 3.00 | 96 |
| 7 | 19 | Male | Unknown | Right | AHR,PHL,PHR | PHL | 1 | 3.00 | 48 |
| 8 | 51 | Female | Hippocampal sclerosis | Right | AHL,AHR,AL,AR,ECL,ECR,PCL,PHR | AHL,PHL | 2 | 3.17 | 150 |
| 9 | 18 | Female | Hippocampal sclerosis | Right | AHL,AHR,AL,ECL,PHL | AHL,PHL | 1 | 3.26 | 23 |
| 10 | 47 | Male | Hippocampal sclerosis | Right | AHL,AHR,AL,AR,ECL,ECR,PHL,PHR | AHR,PHR | 1 | 3.49 | 62 |
| 11 | 37 | Male | Focal cortical dysplasia | Right | AL,AR,HL,HR,FR | FR | 1 | 3.75 | 11 |
| 12 | 24 | Female | Xanthoastrocytoma WHO II | Right | AHL,AL,ECL,LR,PHL,PHR | AHR,LR | 2 | 3.83 | 57 |
| 13 | 39 | Male | Gliosis | Right | AHL,AHR,AL,AR,ECL,ECR,PHL,PHR | AHR,PHR | 1 | 4.50 | 73 |

**Table S1**. Subject characteristics. AH: hippocampal head; PH: hippocampal body; EC: entorhinal cortex; A: amygdala; LL, LR, PL: lesions, AIR: insular gyrus right; FR: frontal right; L: left; R: right; SOZ: seizure onset zone.
